# Amplification-free long read sequencing reveals unforeseen CRISPR-Cas9 off-target activity

**DOI:** 10.1101/2020.02.09.940486

**Authors:** Ida Höijer, Josefin Johansson, Sanna Gudmundsson, Chen-Shan Chin, Ignas Bunikis, Susana Häggqvist, Anastasia Emmanouilidou, Maria Wilbe, Marcel den Hoed, Marie-Louise Bondeson, Lars Feuk, Ulf Gyllensten, Adam Ameur

**Affiliations:** Science for Life Laboratory, Department of Immunology, Genetics and Pathology, Uppsala University, Sweden; Program in Medical and Population Genetics, Broad Institute of Massachusetts Institute of Technology and Harvard, Cambridge, MA, USA; Division of Genetics and Genomics, Boston Children’s Hospital, Boston, MA, USA; Foundation for Biological Data Science; The Beijer laboratory and Department of Immunology, Genetics and Pathology, Uppsala University, Sweden; Department of Epidemiology and Preventive Medicine, Monash University, Melbourne, Australia

**Keywords:** CRISPR-Cas9, on-target, off-target, gRNA, long read sequencing, single molecule sequencing, PacBio sequencing, nanopore sequencing, SMRT-OTS, Nano-OTS

## Abstract

A much-debated concern about CRISPR-Cas9 genome editing is that unspecific guide RNA (gRNA) binding may induce off-target mutations. However, accurate prediction of CRISPR-Cas9 off-target sites and activity is challenging. Here we present SMRT-OTS and Nano-OTS, two amplification-free long-read sequencing protocols for detection of gRNA driven digestion of genomic DNA by Cas9. The methods were assessed using the human cell line HEK293, which was first re-sequenced at 18x coverage using highly accurate (HiFi) SMRT reads to get a detailed view of all on- and off-target binding regions. We then applied SMRT-OTS and Nano-OTS to investigate the specificity of three different gRNAs, resulting in a set of 55 high-confidence gRNA binding sites identified by both methods. Twenty-five (45%) of these sites were not reported by off-target prediction software, either because they contained four or more single nucleotide mismatches or insertion/deletion mismatches, as compared with the human reference. We further discovered that a heterozygous SNP can cause allele-specific gRNA binding. Finally, by performing a *de novo* genome assembly of the HiFi reads, we were able to re-discover 98.7% of the gRNA binding sites without any prior information about the human reference genome. This suggests that CRISPR-Cas9 off-target sites can be efficiently mapped also in organisms where the genome sequence is unknown. In conclusion, the amplification-free sequencing protocols revealed many gRNA binding sites *in vitro* that would be difficult to predict based on gRNA sequence alignment to a reference. Nevertheless, it is still unknown whether *in vivo* off-target editing would occur at these sites.

## Introduction

The CRISPR-Cas9 system is one of the most important breakthroughs in modern biotechnology, as it has provided the possibility to modify the DNA inside living cells in a targeted, easy and cost-efficient manner. CRISPR-Cas9 genome editing was first demonstrated in 2013^1-4^ and has since become an instrumental tool in biomedical research and in bioengineering^5^. CRISPR-Cas9 also shows great promise for clinical use^6^, even though the ethical aspects of human genome editing require careful consideration^7, 8^. A major reason for caution is that the CRISPR-Cas9 system can induce mutations at locations other than the targeted site^9-11^. Such “off-target” mutations have the potential to disrupt the function or regulation of genes in an unpredictive manner, and consequently they are a serious concern for CRISPR-Cas9 applications in the medical field^12^. Development of more efficient and precise genome editing tools systems such as CRISPR-Cas12a^13^ or prime-editing^14^ could help alleviate some of the off-target concerns. But even with these new tools, off-target mutations cannot be excluded, in particular in cases where the DNA sequence of the cells subjected to genome editing is not completely known.

In any CRISPR-Cas9 genome editing experiment, it is crucial to design a guide RNA (gRNA) that specifically binds to the target of interest, and not to any unintended genomic loci. This gRNA will direct the Cas9 endonuclease to its target, after which Cas9 cleaves the DNA molecule, introducing a double stranded break. The DNA is then repaired by non-homologous end joining (NHEJ), and small insertion and deletion mutations are typically introduced during this repair step^15^. In general, Cas9 cleaves its intended target reliably, but off-target mutations can be introduced if the gRNA also binds at other locations. Another potential side effect of CRISPR-Cas9 editing is that larger structural variations, e.g. insertions and deletions of several hundred base pairs, may be introduced during the DNA-repair process by NHEJ. Such large structural variants (SVs) have been detected at the on-target site^16^, but they have not yet been shown to occur at off-target sites. Although there have been conflicting reports on the abundance and consequences of unintended mutations^16-19^, there is a consensus that off-target sites should be screened for when designing a gRNA, to increase the chances of a successful and specific genome editing^12^.

Guide RNAs are typically designed by computational tools that compare the gRNA sequence to a reference genome and predict the binding affinity both to the on-target sequence as well as to potential off-targets^20-22^. Although intuitively helpful, these tools can yield false positive or negative results due to the difficulty to exactly model gRNA-DNA binding affinity in an algorithm. Furthermore, the DNA sequence in the cells being investigated can differ substantially from the reference genome used in the computational modeling, potentially resulting in even more false predictions. In recent years, *in vitro* based assays such as Digenome-seq^23^ and CIRCLE-Seq^24^ have been developed that allows to experimental detection of gRNA binding sites in a particular DNA sample. However, since these methods are based on PCR amplification and short-read sequencing, they have inherent limitations when it comes to detection of gRNA binding in repetitive, low complexity, or AT/GC-rich regions. These issues can be improved by long-read single molecule sequencing technologies. At present, Pacific Biosciences (PacBio) and Oxford Nanopore Technologies (ONT) are the two main providers of long-read sequencing, and it is now widely accepted that these technologies have a superior ability, as compared to short-read sequencing, to resolve SVs as well as other complex regions in the human genome^25-29^.

Here we propose two new methods for accurate *in vitro* detection of gRNA binding and Cas9 cleavage, and we denote these “off-target sequencing” (OTS). The methods are based on PacBio’s single molecule real-time sequencing (SMRT-OTS) and ONT’s nanopore sequencing (Nano-OTS). By introducing these two protocols, rather than just one, we have an alternative amplification-free method to employ for orthogonal validation of our findings. The SMRT-OTS and Nano-OTS methods were evaluated using DNA from the human HEK293 cell line. Importantly, the HEK293 cells were whole genome sequenced to high coverage using long and accurate SMRT sequencing reads, i.e. high-fidelity (HiFi) reads^29^, to get the best possible view of the genomic DNA to which the gRNA binds.

## Results

### Two new amplification-free protocols for off-target sequencing

We developed two methods for gRNA off-target sequencing (OTS) (see **Figure 1A-B**). SMRT-OTS is based on PacBio’s SMRT sequencing and produces highly accurate circular consensus reads, which can be used to detect the exact gRNA binding sequence as well as genetic variants in on- and off-target regions. Nano-OTS is based on ONT’s nanopore sequencing and allows for rapid identification of gRNA binding sites but with lower per-read accuracy. Our methods are inspired by previously proposed assays where a single gRNA was used to perform targeted enrichment. SMRT-OTS is a modified version of a protocol we previously applied for detection of repeat expansions in human cell lines and blood samples^30, 31^, while Nano-OTS is adapted from a targeted sequencing assay^32^ used for detection of unknown fusion gene partners^33^. In addition to wet lab assays, we developed a computational method that can be used to identify gRNA binding sites a single base pair resolution, both from high-quality SMRT reads and from lower quality nanopore reads (see **Figure 1C**). In the analysis, candidate gRNA binding sites are found from specific patterns in the alignment where several reads start or end at the exact same position. Because the reads from our OTS assays originate from randomly sheared DNA fragments with varying start and end positions, such patterns are highly unlikely to arise from background reads that have not been cleaved by Cas9. Pairwise alignments are then performed between all gRNA sequences and all predicted Cas9 cleavage regions to determine which gRNA is bound to which target. In this step, false positive peaks with little or no resemblance to any gRNA sequence are removed.

**Figure 1.**
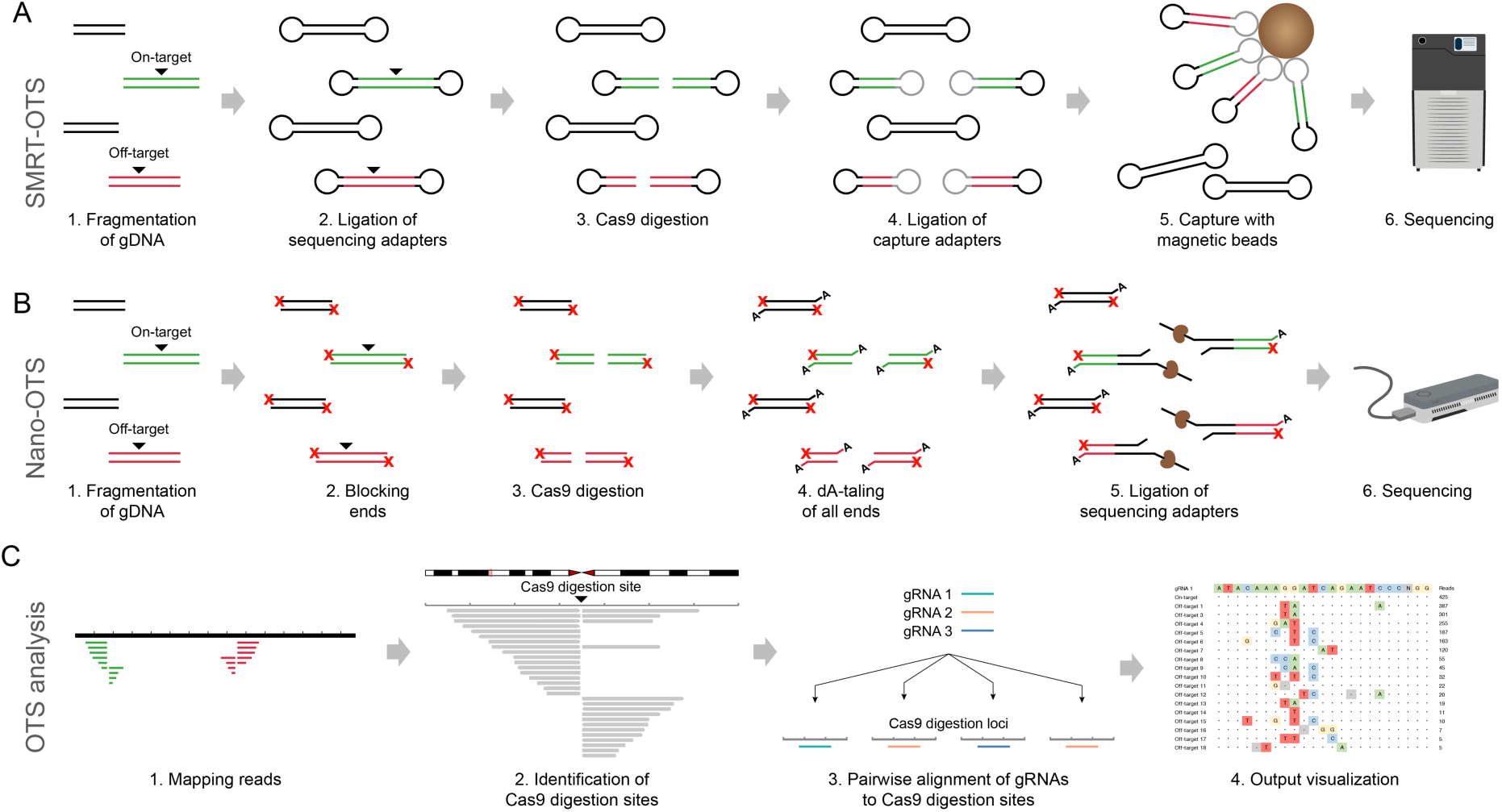
The SMRT-OTS and Nano-OTS methods for gRNA binding detection. **A)** SMRT-OTS starts from a DNA sample that has been fragmented by random shearing and can be applied either to a single gRNA or to a pool of several gRNAs. The DNA fragments will either contain an intended on-target site (green), an off-target (red) or no gRNA binding site (black) (1). After fragmentation, a SMRT bell library is prepared by ligating sequencing adapters at both ends (2). The SMRT bells containing gRNA binding sites are then cleaved by Cas9 (3), after which capture adapters (gray) are ligated to the cleaved molecules (4). Finally, magnetic beads are used to capture SMRT bells containing the capture adapter (5). This gives an enrichment of SMRT bells cleaved by Cas9 and the enriched library is sequenced on the PacBio Sequel system (6). **B)** Nano-OTS, just like SMRT-OTS, starts from a randomly fragmented DNA sample and can either be applied to a single gRNA or to a multiplexed pool of gRNAs (1). All ends of the fragmented DNA molecules are dephosphorylated to block the ends from adapter ligation (2). The DNA fragments containing gRNA binding sites are cleaved by Cas9 (3) and all 3’ ends are dA-tailed (4). Sequencing adapters are then ligated to Cas9 cleaved and dA-tailed ends (5) and these molecules are sequenced on the ONT MinION system (6). **C)** The same computational approach is used both for SMRT-OTS and Nano-OTS. Reads are aligned to a reference genome, and this gives rise to specific patterns at gRNA on- and off-target binding sites with multiple reads starting at the same position (1). An in-house developed script searches for such patterns in the alignment file and reports all detected targets in a bed file (2). The reference sequence is extracted for all the targets and aligned to the gRNA sequences. Only regions with sufficient similarity to a gRNA sequence are kept (3). The results are reported in an output file that contains the Cas9 cleavage position, the sequence alignment of the gRNA to the reference, and the peak height from the OTS sequencing (4).

### Detection of gRNA binding in human HEK293 cells using SMRT-OTS

DNA from the human embryonic kidney cell line HEK293 was used to evaluate the OTS protocols. As a baseline for our experiments, a comprehensive genome map of the HEK293 cells was generated by HiFi SMRT sequencing^29^, resulting in 18x whole genome coverage with >Q20 reads of an average read length of 15 kb. Because of their length and accuracy, the HiFi reads are ideal both for detection of single nucleotide variants (SNVs) as well as larger SVs^29^. After having determined the HEK293 genome sequence, we performed a multiplexed SMRT-OTS run with three guide RNAs, designed to target an intron of *ATXN10*, and early exons of *MMP14* and *NEK1*. These three gRNAs have all been used in previous experiments, by us and others (see Methods). Sequencing was performed on a Sequel 1M SMRT cell, resulting in a total of 57,644 reads with an average read length of 4.0 kb. All three gRNA on-target sites were successfully detected, along with 42 off-targets for *ATXN10*, 27 off-targets for *MMP14*, and three off-targets for *NEK1* (see **supplementary Data S1**). The on-target alignment peaks for the three gRNAs, as well as examples of off-target peaks with at least three mismatches to the HEK293 genome, are shown in **Figure 2**.

**Figure 2.**
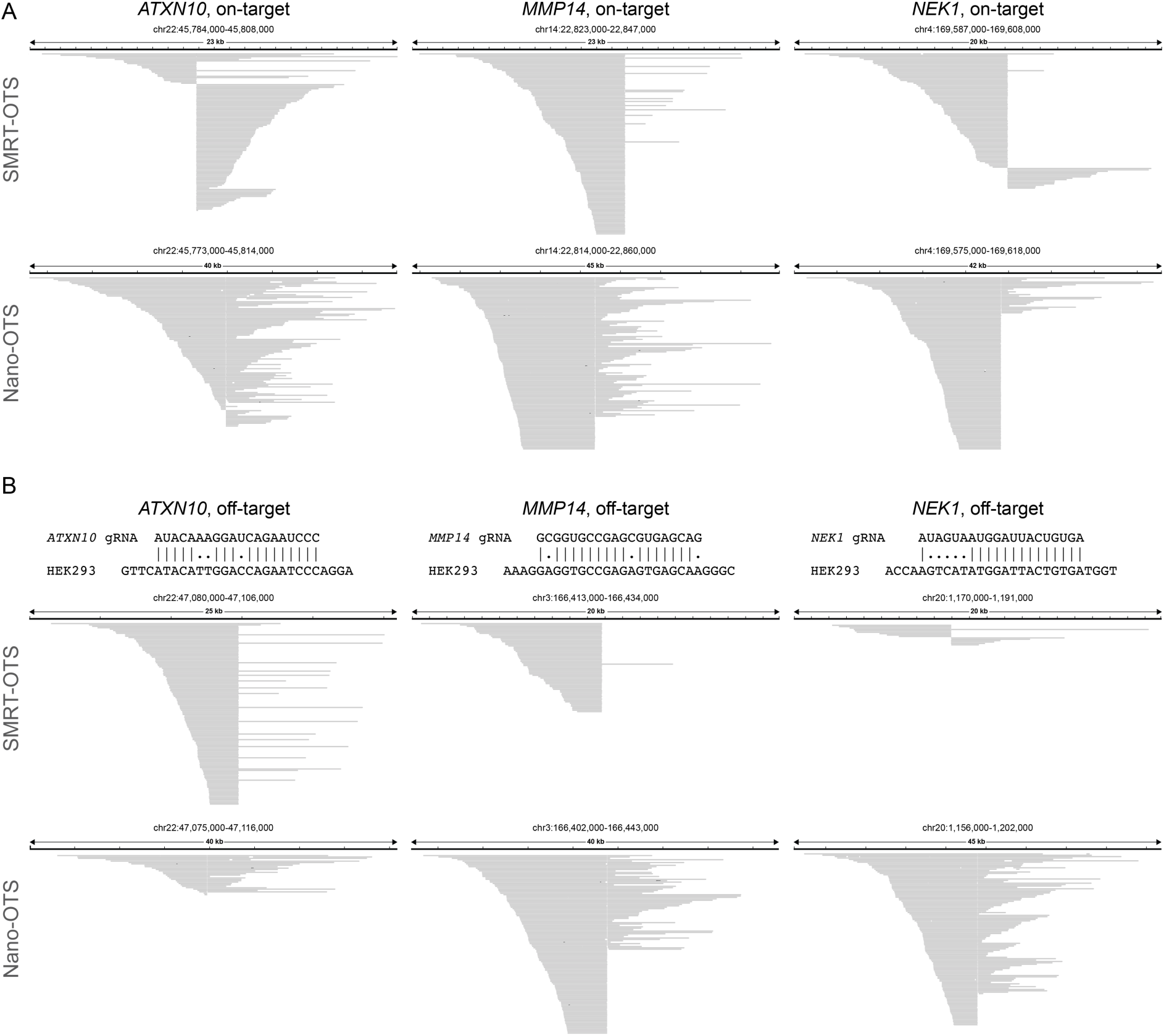
Examples of detected on- and off-target peaks in HEK293 DNA. **A)** IGV alignments^46^ showing the read distributions for SMRT-OTS (top) and Nano-OTS (bottom) in 20-50 kb windows spanning the *ATXN10, MMP14* and *NEK1* on-target sites. The Cas9 cleavage sites are clearly visible as sharp vertical lines where the alignments start or end. **B)** Examples of off-target alignment peaks for *ATXN10* (left), *MMP14* (middle) and *NEK1* (right). At the top are sequence alignments between the gRNA sequence and the HEK293 genome at the off-target site. There are three single nucleotide mismatches at the *ATXN10* and *MMP14* off-target sites, and five single nucleotide mismatches at the *NEK1* off-target site. All three off-target sites are visible in both in the SMRT-OTS and Nano-OTS alignment data.

### Validation of gRNA binding in HEK293 cells using Nano-OTS

To validate our results and to examine the reproducibility of our sequencing protocols, we performed a Nano-OTS run using the same HEK293 DNA and the same three gRNAs. 185,145 reads of average length 7.5 kb were generated on one MinION flow cell. Fifty-four, 30 and 50 off-targets were found for *ATXN10, MMP14*, and *NEK1*, respectively (**supplementary Data S2**). Due to a higher error rate in the Nano-OTS reads, the gRNA binding sites are sometimes predicted within an 10-20 bp interval instead of at exact base pair resolution. Fifty-five gRNA binding sites overlapped between the two methods, while 20 were found only by SMRT-OTS and 82 only by Nano-OTS (see **Figure 3A** and **supplementary Data S3**). There is a positive correlation in peak heights when comparing the SMRT-OTS and Nano-OTS results, even though the signals can differ substantially for individual sites (**Figure 3B**). The discrepancy in results can partly be explained by the higher coverage in the Nano-OTS run, but there might also be differences in Cas9 cleavage efficiency between the two methods.

**Figure 3.**
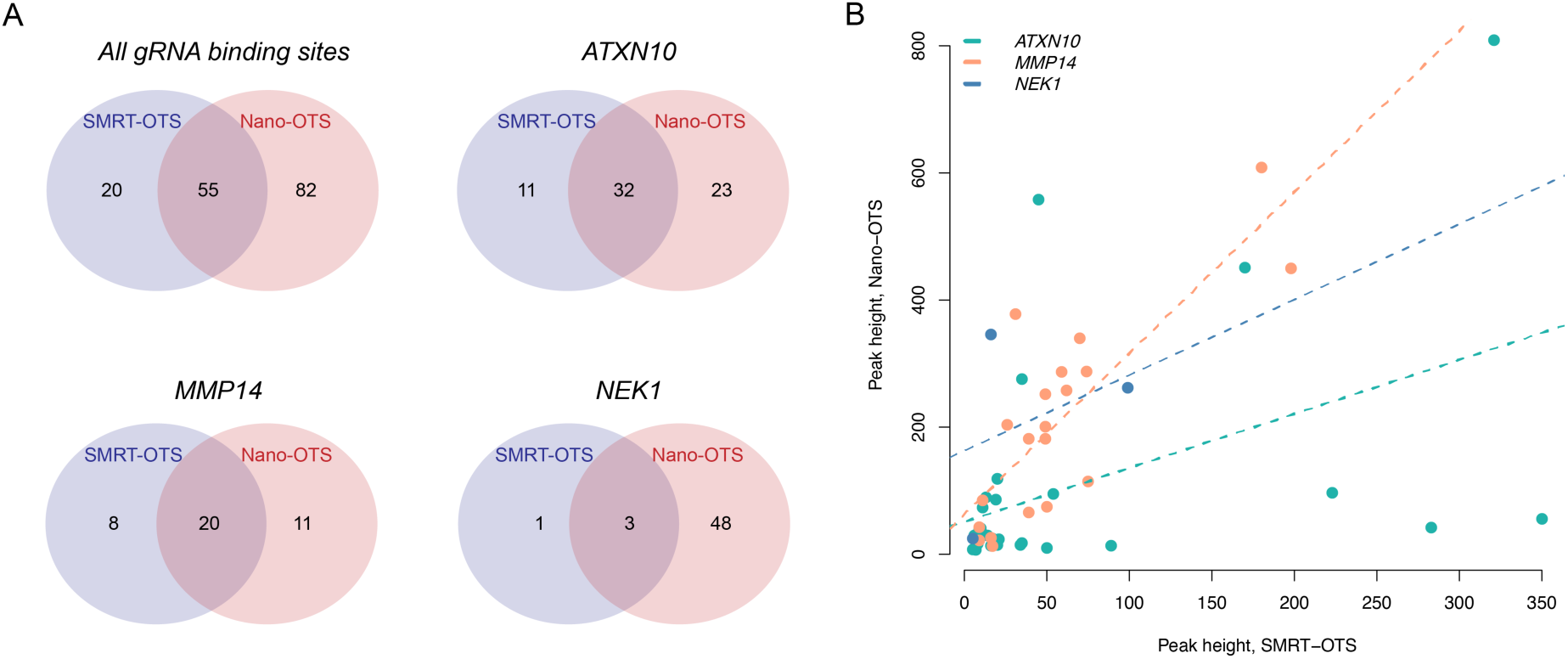
Comparison of results from SMRT-OTS and Nano-OTS. **A)** Venn diagrams showing the overlap between gRNA predictions for SMRT-OTS and Nano-OTS for all three gRNAs combined (top left panel) and each individual gRNA (the three other panels). **B)** Correlation between peak heights for SMRT-OTS and Nano-OTS for the 55 gRNA binding predictions detected by both methods. Binding sites for *ATXN10, MMP14* and *NEK1* gRNAs are plotted in different colors, and the dotted lines show the regression lines. Even though there are large differences between SMRT-OTS and Nano-OTS peaks in some cases, there is a positive association when considering all data points.

### Guide RNAs may bind to off-targets despite high sequence dissimilarity

As SMRT-OTS and Nano-OTS are two orthogonal methods, we considered the intersection of their results to be a high-confidence set of targets predicted to be cleaved by Cas9 in the HEK293 cells. We used the term “OTS-targets” for this dataset, which, as mentioned above, consists of 55 gRNA binding sites, and is presented in detail in **Figure 4**. For comparison, we used the latest release of CHOPCHOP^22^ to predict gRNA binding *in silico*. A total of 82 binding sites were reported by CHOPCHOP when allowing for up to three single nucleotide mismatches (**supplementary Data S4**). Of these *in silico* predictions, as many as 45 (55%) were not detected by OTS-SMRT or OTS-Nano. These could either be sequences not bound by a gRNA despite high similarity, sites bound by a gRNA but not cleaved by Cas9, or a combination of both. Conversely, 25 (45%) of our OTS-targets were not reported by CHOPCHOP (see **supplementary Table S1**). Among these, 18 OTS-targets had at least four single nucleotide mismatches to the gRNA sequence, and seven OTS-targets contained insertion/deletion mismatches to the gRNA sequence. Three of the OTS-targets have a mismatch in the PAM sequence (NGG), but for all those cases a canonical PAM sequence can be found at the subsequent position.

**Figure 4.**
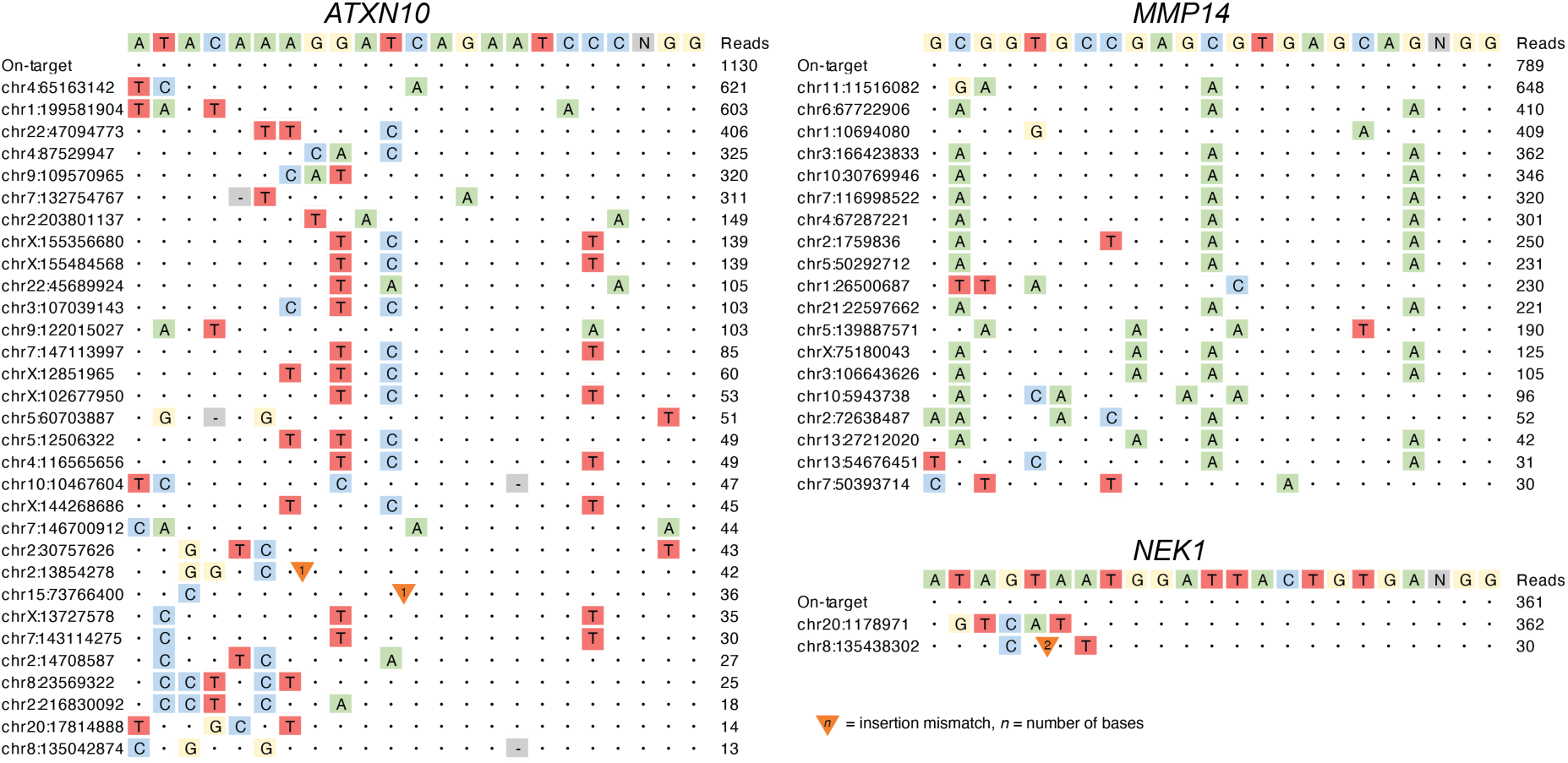
High confidence gRNA binding sites in HEK293 cells. Visualization of the 55 gRNA binding sites for *ATXN10, MMP14* and *NEK1* that were detected both by SMRT-OTS and Nano-OTS. The letters at the top show the gRNA sequence and the bases (NGG) that corresponds to the PAM site. Below are the on-target as well as all off-target sequences detected by the OTS methods. Colored letters correspond to single nucleotide mismatches in the HEK293 genome as compared with the gRNA sequence. Triangles and hyphens (-) are used to mark sites where nucleotides need to be inserted or deleted, respectively, in the HEK293 genome order to match the gRNA sequence. The column to the right contains the combined read count from the SMRT-OTS and Nano-OTS assays, for each gRNA binding site.

### A single nucleotide polymorphism can induce allele-specific gRNA binding

The HiFi data for HEK293 allowed us to identify and phase genetic variants across the entire genome. Based on this information, we could investigate allelic biases in gRNA binding. One allele-specific binding event was found, at an off-target site for *ATXN10*. At this site HEK293 was reported heterozygous for the T/C SNV rs7861875 (see **Figure 5A**). The HiFi data further revealed a haplotype with several additional SNPs in the region, all of them linked to the reference allele of rs7861875 (T). The rs7861875 T allele and associated SNV haplotype is present in six of 23 HiFi reads (26%), and the deviation from 50% as well as elevated coverage in the HiFi data suggests that this locus may be duplicated in HEK293 cells (**supplementary Figure S1**). In the SMRT-OTS data, 101 of the 106 reads (95%) contain the alternative allele, and only five reads (5%) carry the T allele and associated SNV haplotype. This is consistent with a preferential gRNA binding to the C allele, which has higher sequence similarity (two mismatches) to the *ATXN10* gRNA as compared with the T allele (three mismatches) (see **Figure 5B**). Rs7861875 has a C allele frequency of 0.20 in gnomAD^34^, suggesting that about 20% of human individuals would be affected by *ATXN10* gRNA off-target binding at this site, while those homozygous for the T allele would be largely unaffected. Although this is just one example, it demonstrates that common genetic variation can cause unintended gRNA binding and that our methods are sensitive enough to identify such events. Only one additional heterozygous SNV was present in a gRNA binding site (*MMP14*; chr2:1759836), but the SMRT-OTS coverage in that region was too low to study allele specific binding.

**Figure 5.**
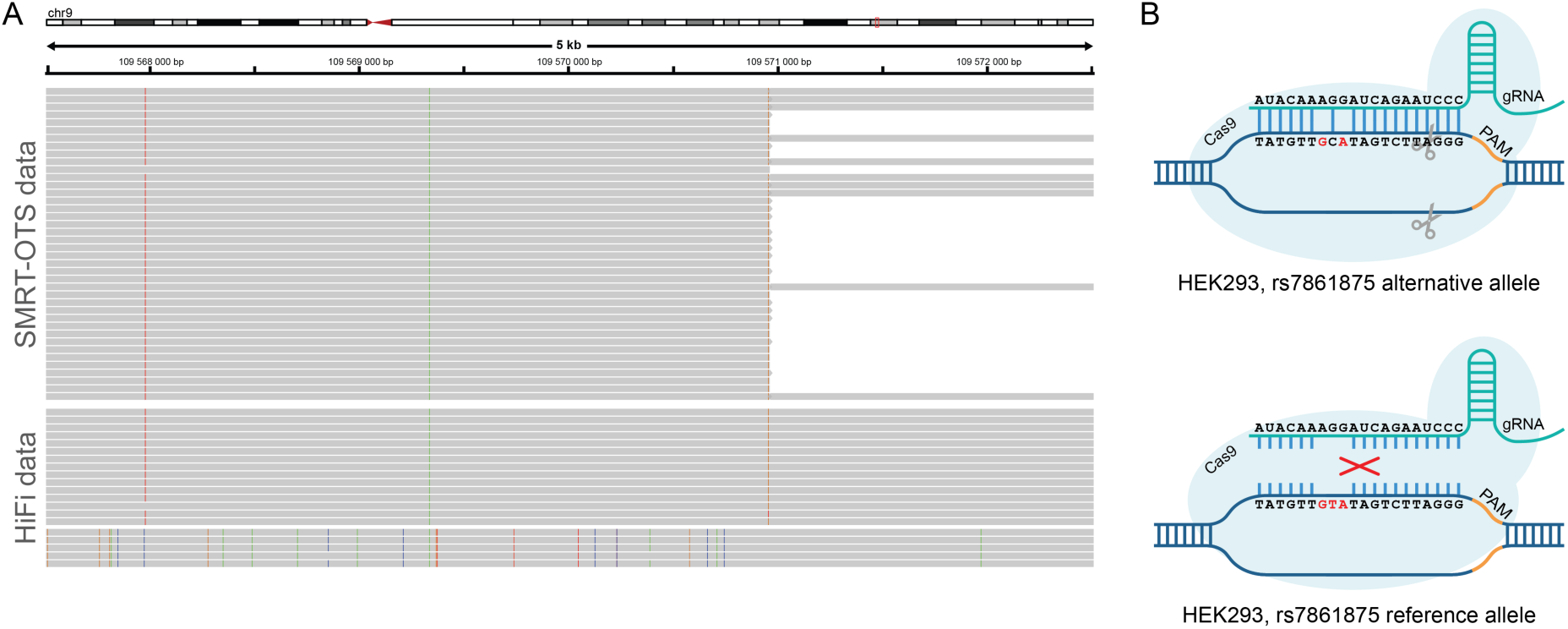
Genetic variation in HEK293 induces allele-specific gRNA off-target binding. **A)** An IGV image showing SMRT-OTS alignments (top) and HEK293 HiFi SMRT sequencing reads (bottom) in a window surrounding a predicted gRNA site for *ATXN10* (chr9:109,570,956; GRCh38 coordinates). The HiFi reads reveal a heterozygous SNP (rs7861875; T>C) in HEK293 within the *ATXN10* gRNA binding site, where the reference allele (T) is linked to the alternative alleles of several other heterozygous SNPs in the window. In the SMRT-OTS data, 101 of the 106 reads (95%) of the reads correspond to the haplotype having the alternative rs7861875 allele (G). **B)** Schematic view of allele specific *ATXN10* gRNA binding at the rs7861875 locus. The CRISPR-Cas9 complex does not bind to the rs7861875 reference allele (T), which has three mismatches to the gRNA sequence (bottom). It binds more efficiently to the alternative allele (C) which has only two mismatches to the *ATXN10* gRNA sequence (top).

### Searching for gRNA binding in “dark matter” regions of the HEK293 genome

Human long-read assemblies have been shown to contain several megabases (Mbs) of novel sequences or alternative haplotypes with high diversity from the GRCh38 reference^35, 36^. We were interested to determine whether any additional off-targets could be detected in such “dark matter” regions of the HEK293 genome. Hence, we first prepared a *de novo* assembly of the HEK293 HiFi data was performed using Peregrine^37^, resulting in a genome size of 2,896 Mb with N50 of 11.2 Mb (**supplementary Table S2**). We next aligned the SMRT-OTS reads to the HEK293 *de novo* assembly and could identify 43, 27, and four gRNA binding sites for *ATXN10, MMP14* and *NEK*, respectively (see **supplementary Data S5**). By comparing the sequences in the binding regions, we concluded that all of these sites were detected among the 75 SMRT-OTS predictions obtained for GRCh38. Even though no additional off-targets were detected in dark matter regions of HEK293, our results suggest that *de novo* assembly in combination with OTS is a powerful method to screen for off-target activity in organisms without a readily available reference genome, since as many as 74 of the 75 (98.7%) of gRNA binding sites found in GRCh38 could be identified in this way.

## Discussion

With long-read single molecule sequencing, global patterns of gRNA activity can be monitored at a new level of resolution. Existing methods are based on Illumina sequencing^23, 24^ and are therefore restricted to short reads that are not suited to interrogate repetitive and highly polymorphic regions. Moreover, Illumina sequencing relies on amplification during library preparation that may lead to biases and uneven sequence coverage^38^. Long-read sequencing, on the other hand, overcomes some of the short-read limitations through unbiased single molecule coverage and average read lengths of several kilobases. Because of the amplification-free nature of the SMRT-OTS and Nano-OTS protocols, each read supporting an on- or off-target site originates from an individual DNA molecule that has been cleaved by Cas9. We therefore hypothesize that the peak heights resulting from the OTS-protocols could be used as a proxy to quantify gRNA binding strength and CRISPR-Cas9 genome editing efficiency. A further advantage of sequencing native DNA molecules is that base modification signals can be detected^39, 40^, thereby paving the way for future studies on how epigenetic events and chromatin structure affect CRISPR gRNA activity^41^.

Although SMRT-OTS and Nano-OTS are both based on amplification-free long-read sequencing, each method has its own unique features. SMRT-OTS has the advantage of producing high quality circular consensus sequences, thereby enabling accurate SNV calling, detection of allele specific gRNA binding and identification of Cas9 cleavage sites at base pair resolution. Nano-OTS on the other hand is a faster protocol, that can be performed within just one or two days, and using a portable, cheaper sequencing instrument. Both SMRT-OTS and Nano-OTS allow multiplexing of several gRNAs in a single run. In this study, we tested up to three gRNAs on PacBio’s Sequel and ONT’s MinION instruments, but it should be possible to increase the degree of multiplexing by an order of magnitude using the higher throughput Sequel II and PromethION systems. Higher order multiplexing could be useful when screening large gRNA panels for optimal candidates in gene knockout experiments, or for post-hoc quantification of off target effects in such studies. One limiting factor, however, is that the protocols require substantial amounts of high-quality input DNA. In our current setup, SMRT-OTS requires ∼10-15 µg of DNA as starting material, and Nano-OTS ∼5-10 µg. We are hopeful these numbers can be reduced by further optimization of the protocols.

A unique aspect of our gRNA binding experiment is that we determined the exact genetic background of the HEK293 cells. For this, we used state-of-the-art high HiFi whole genome sequencing^29^. The HEK293 HiFi data, coupled with results from the OTS-assays, gives us a more detailed view of gRNA on- and off-target activity in human DNA than ever before. In fact, we were able to detect a vast majority of the gRNA binding sites *de novo*, without making use of the existing human reference genome. One intriguing finding was preferential binding of the *ATXN10* gRNA to the alternative allele at rs7861875. Although one should be careful to draw general conclusions from a single example, this result suggests that SNVs can induce unexpected off-target activity and that individual level genetic variation should be taken into consideration when designing gRNAs for medical purposes. Computational strategies that take into account SNVs in gRNA binding prediction already exist^42, 43^, but those rely on databases that does not contain all variants that any individual carries. Our results further demonstrate that gRNAs can bind to genomic DNA despite having three or more single nucleotide mismatches, or even insertion or deletion mismatches. Since binding sites with high sequence divergence are difficult to predict using computational tools, we argue that *in vitro* tools like the ones presented here are needed to accurately determine where a gRNA binds in a particular DNA sample. However, more experiments are required to understand whether off-targets confirmed *in vitro* also result in off-targets mutations *in vivo*, or whether there are protective mechanisms *in vivo* that would prevent off-target CRISPR/Cas9 genome editing.

In summary, with SMRT-OTS and Nano-OTS, we provide new and efficient tools to evaluate and improve gRNA design, as well as to optimize CRISPR protocols. Coupled with high accuracy long-read whole genome sequencing, we believe these methods will enable us to better understand the mechanisms of gRNA binding and, hopefully, also to prevent negative effects of off-target and unintended mutations in future CRISPR-Cas9 experiments.

## Acknowledgements

We thank the R&D teams at Pacific Biosciences and Oxford Nanopore Technologies for their assistance. SMRT sequencing was performed by the SciLifeLab National Genomics Infrastructure (NGI) in Uppsala, Sweden. Computations were performed on resources provided by SNIC through Uppsala Multidisciplinary Center for Advanced Computational Science (UPPMAX) under Project b2017186.

## Materials and Methods

### Samples

Genomic DNA from the HEK293 cell line was purchased from GenScript (https://www.genscript.com).xs

### Whole genome HiFi SMRT sequencing of HEK293 on Sequel II

To generate a HiFi library, genomic DNA was sheared using the Megaruptor 2 (Diagenode) with a long hydropore and a 20 kb shearing protocol. Size distribution of the sheared DNA was characterized on the Femto Pulse system (Agilent Technologies) using the Genomic DNA 165 kb Kit. Sequencing libraries were constructed using the protocol “Preparing HiFi SMRTbell Libraries using SMRTbell Express Template Prep Kit 2.0” from PacBio. SMRTbells were size selected using 0.75% agarose 1-18kb protocol on SageELF (Sage Science) according to the HiFi SMRTbell library protocol. Size selected SMRTbells were examined on the Femto Pulse system (Agilent Technologies) using the Genomic DNA 165 kb Kit. Library fraction of 15 kb and 17 kb were selected for sequencing. Sequencing was performed on two SMRT cells using the Sequel II system and the 2.0 sequencing and binding chemistry, with 2 hours pre-extension and 30 hours movie time.

### Guide RNAs

All three gRNAs used in this study have all been used in previous experiments, and they were purchased from Integrated DNA Technologies. The *ATXN10* gRNA (AUACAAAGGAUCAGAAUCCC) is the same as that used in our previous experiments on amplification-free PacBio sequencing of repeat expansions in the human genome^24, 31^. The *MMP14* gRNA (GCGGUGCCGAGCGUGAGCAG) has been used by us in genome editing experiments. The *NEK1* gRNA (AUAGUAAUGGAUUACUGUGA) has been used in genome editing experiments by Horizon Discovery, and a *NEK1* edited HAP1 cell line can be ordered from their website (https://horizondiscovery.com).

### SMRT-OTS: Off-target sequencing using PacBio’s SMRT sequencing

SMRT-OTS libraries were prepared in a similar manner described by Tsai et al^30^, with modifications. Genomic DNA was sheared to 8 kb fragments using Megaruptor 2 (Diagenode). Standard SMRTbell libraries were prepared using Template Preparation Kit 1.0 (Pacific Biosciences) according to manufacturer’s instructions. An extra exonuclease treatment, using Exonuclease I (New England Biolabs) and Lambda exonuclease (New England Biolabs) was added at the end of the library preparation. The final SMRTbell library was size selected using the Blue Pippin system (Sage Science) with a cut-off at 4 kb. The crRNA and tracrRNA with Alt-R modification (Integrated DNA Technologies) were annealed in a 1:1 ratio to form gRNA that was used in the Cas9 (New England Biolabs) digestion of the SMRTbell libraries. Cas9 and gRNA in presence of buffer were incubated at 37°C for 10 minutes, before heparin was added and the mixture was incubated for an additional 3 minutes at 37°C. 1 µg of SMRTbell library was then added and incubated for 1 hour at 37°C. EDTA was then added to terminate the reaction and the SMRTbell library was subjected to PB AMPure bead (Pacific Biosciences) purification. Hairpinned capture adapters with a polyA-stretch (5’-ATCTCTCTCTTAAAAAAAAAAAAAAAAAAAAAAATTGAGAGAGAT-3’) were ligated, overnight at 16°C, to the Cas9 digested SMRTbell molecules using T4 DNA ligase (Thermo Fischer Scientific) forming asymmetrical SMRTbell libraries. The asymmetrical SMRTbell library were subjected to exonuclease III and VII (Pacific Biosciences) at 37C for 1 hr. MagBeads (Pacific Biosciences) were used to enrich for asymmetric SMRTbell molecules by binding to the capture hairpin-adapters. The asymmetric SMRTbell molecules/MagBead complex was incubated under rotation at 4°C for 2 hours in MagBead Binding buffer v2 (Pacific Biosciences) three times. Finally, the enriched asymmetric SMRTbells were eluted in Elution buffer (Pacific Biosciences) for 10 min at 50°C. The asymmetric SMRTbell molecules were prepared for SMRT sequencing by primer annealing with standard PacBio sequencing primer lacking the polyA sequence for 1 hour at 20°C. Sequel DNA polymerase 3.0 was bound to the template/primer complex for 4 hours at 30°C. Sequencing was performed on the PacBio Sequel system using one 1M SMRT cell, Sequel Sequencing kit 3.0 and a 600 min movie time. Asymmetric SMRTbell template sequencing data was subjected to a customized analysis pipeline for capture and conventional hairpin-adapter recognition for separating subreads. Subsequently, the CCS tool in SMRT analysis was used to create circular consensus sequencing reads from the subreads.

### Nano-OTS: Off-target sequencing using ONT’s nanopore sequencing

Genomic DNA was sheared to 20 kb fragments using Megaruptor 2 (Diagenode) and size selected using the BluePippin system (Sage Science) with a cut-off at 10 kb. 3-4 µg of sheared and size-selected DNA was prepared using the Cas9-mediated PCR-free protocol provided by Oxford Nanopore technologies with minor modifications. The crRNA and tracrRNA with Alt-R modification (Integrated DNA Technologies) were annealed in Duplex buffer (Integrated DNA Technologies) at 95°C for min and were then allowed to cool down to room temperature. Ribonucleoproteins (RNPs) were formed by combining the annealed gRNA, HiFi Cas9 (Integrated DNA Technologies) and 1x NEB CutSmart buffer (New England Biolabs) and incubated at room temperature for 30 min. The fragmented and size-selected DNA was dephosphorylated to block all ends from ligation of adapters in a downstream adapter ligation step. Subsequently, the DNA molecules were digested by Cas9 using the previously prepared RNPs and the newly cleaved ends were dA-tailed to enable adapter ligation. The library preparation was completed by ligation of adapters from the SQK-LSK109 kit (Oxford Nanopore Technologies) and cleaned up with AMPure XP beads (Beckman Coulter) before preparation for sequencing. Sequencing was performed using the MinION system (Oxford Nanopore Technologies) with a R9.5.1 flow cell and Guppy v3.3.3 was used for base calling.

### Alignment of reads and detection of off-target gRNA binding sites

The reads from SMRT-OTS and Nano-OTS were aligned to GRCh38 using minimap2^44^, after which gRNA binding sites were predicted using v1.8 of the Insider software (https://github.com/UppsalaGenomeCenter/InSiDeR). For each predicted gRNA binding site, the corresponding sequence from GRCh38 was extracted in a +-40 bp window surrounding the Cas9 cleavage site. All sequences containing gaps (N’s) were filtered out since we were only interested in detection of gRNA binding event in high-quality regions of the human genome. For the remaining sequences, we performed global alignment against all gRNA sequences using v6.6.0 of EMBOSS-Needle with default settings^45^. Only sequences with containing an alignment score of >55 to a certain gRNA were considered positive binding sites.

### De novo assembly of HEK293 HiFi SMRT sequencing data

Data from two HiFi Sequel II SMRTcells were assembled with Peregrine build 0.1.6.0, using a docker image on an AWS r5d.12xlarge instance. The command options are available in Supplementary Information.

