## Supplementary Information for "Amplification-free long read sequencing reveals unforeseen CRISPR-Cas9 off-target activity"

### Supplementary Tables

**Supplementary Table S1.** OTS sites found both by SMRT-OTS and Nano-OTS, but not detected by CHOPCHOP.

| chr | position | gRNA | DNA binding sequence | mm | indels | SMRT | Nano |
|---|---|---|---|---|---|---|---|
| chr1 | 199581904 | atxn10 | TAATAAAGGATCAGAATACCAGG | 4 | 0 | 45 | 558 |
| chr7 | 132754767 | atxn10 | ATAC-TAGGATCAAAATCCCAGG | 2 | 1 | 35 | 276 |
| chr2 | 13854278 | atxn10 | CCAGGGATTCTGATCCATGTCCAT | 3 | 1 | 17 | 25 |
| chr7 | 146700912 | atxn10 | CAACAAAGGATAAGAATCCCAAG | 4 | 0 | 14 | 30 |
| chr2 | 30757626 | atxn10 | ATGCTCAGGATCAGAATCCCATG | 4 | 0 | 12 | 31 |
| chr5 | 60703887 | atxn10 | CATGGGATTCTGATCCTCT-TCT | 3 | 1 | 10 | 41 |
| chr10 | 104676047 | atxn10 | CCAGGGA-TCTGATGCTTTGTGA | 3 | 1 | 9 | 38 |
| chr2 | 14708587 | atxn10 | CCAGGGATTCTGTTCCTGAGTGT | 4 | 0 | 8 | 19 |
| chr8 | 23569322 | atxn10 | ACCTACTGGATCAGAATCCCTGG | 5 | 0 | 8 | 17 |
| chr20 | 17814888 | atxn10 | TTAGCATGGATCAGAATCCCGGG | 4 | 0 | 7 | 7 |
| chr2 | 216830092 | atxn10 | ACCTACAGAATCAGAATCCCTGG | 5 | 0 | 7 | 11 |
| chr15 | 73766400 | atxn10 | ATCCAAAGGATACAGAATCCCAGG | 1 | 1 | 6 | 30 |
| chr8 | 135042874 | atxn10 | CCAGGGA-TCTGATCCTCTGCAG | 3 | 1 | 5 | 8 |
| chr5 | 139887571 | mmp14 | GCAGTGCCAAGCATGAGTAGTGG | 4 | 0 | 75 | 115 |
| chrX | 75180043 | mmp14 | GAGGTGCCAAGAGTGAGCAAGGG | 4 | 0 | 50 | 75 |
| chr2 | 1759836 | mmp14 | GAGGTGCTGAGAGTGAGCAAGGG | 4 | 0 | 49 | 201 |
| chr3 | 106643626 | mmp14 | CCCTTGCTCACTCTTGGCACCTC | 4 | 0 | 39 | 66 |
| chr1 | 26500687 | mmp14 | CCCCTGCTCAGGCTCGGCTCAAC | 4 | 0 | 26 | 204 |
| chr7 | 50393714 | mmp14 | CCTGTGCTGAGCGTAAGCAGTGG | 4 | 0 | 17 | 13 |
| chr13 | 27212020 | mmp14 | CCCTTGCTCACTCTTGGCACCTC | 4 | 0 | 16 | 26 |
| chr10 | 5943738 | mmp14 | CCCCTGCTCATGTTCGGTGCCTC | 5 | 0 | 11 | 85 |
| chr13 | 54676451 | mmp14 | CCCTTGCTCACTCTCGGCGCCGA | 4 | 0 | 9 | 22 |
| chr2 | 72638487 | mmp14 | AAGGTACCGAGAGTGAACAGAGG | 5 | 0 | 9 | 43 |
| chr20 | 1178970 | nek1 | CCATCACAGTAATCCATATGACT | 5 | 0 | 16 | 346 |
| chr8 | 135438302 | nek1 | CCTTCACAGTAATCCAATGAAGTAT | 2 | 2 | 5 | 25 |

**Supplementary Table S2.** Results of *de novo* assembly of HEK293 HiFi data

| | **HEK293 *de novo* assembly statistics** |
|---|---|
| Total contigs | 2,110 |
| Total bases | 2,896,467,641 |
| Max contig length | 47,094,793 |
| N50 | 11,221,406 |
| N90 | 1,874,419 |
| N95 | 724,820 |

#### Supplementary Figures

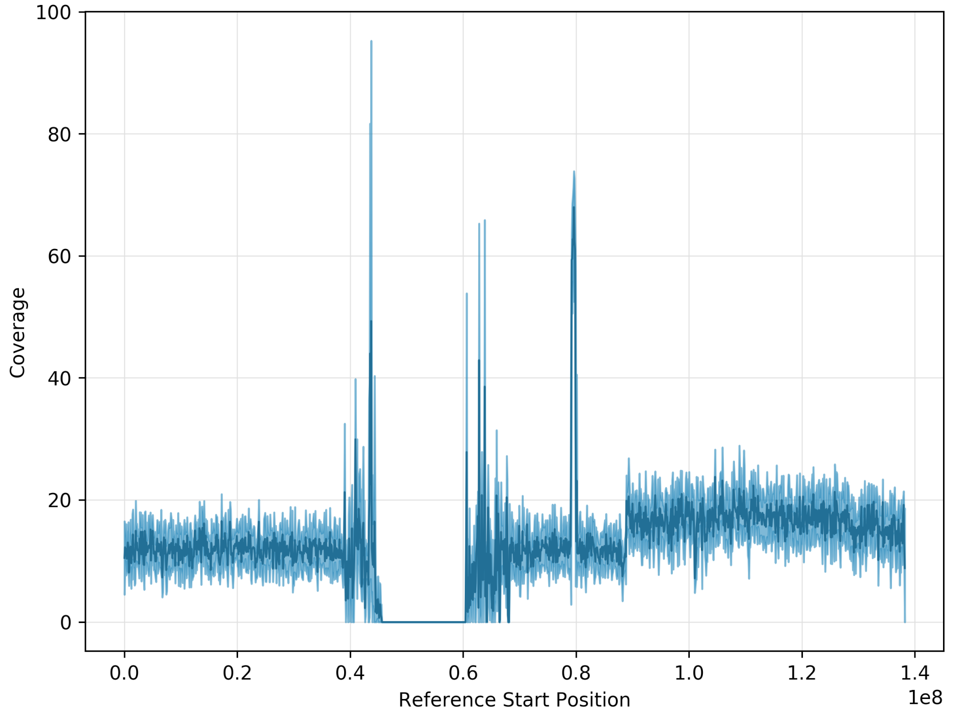


**Supplementary Figure S1.** Coverage profile of HEK293 HiFi data on chromosome 9. The SNP rs7861875 (chr9:109,570,956) is located near coordinate 1.1 in a part of the chromosome that has elevated coverage as compared to the rest of the chromosome.

#### Supplementary Information

SMRTcell data was assembled with Peregrine build 0.1.6.0 docker image with the following command options on an AWS r5d.12xlarge instance:

*docker run -it -v /wd:/wd --user $(id -u):$(id -g) cschin/peregrine:0.1.6.0 asm /wd/seqdata.lst 24 24 24 24 24 24 24 24 24 --with-consensus --shimmer-r 3 --best_n_ovlp 8 --output /wd/asm*

The file /wd/seqdata.lst contains the file path to the input file:

*/wd/data/m64077_191107_133924.Q20.fasta.gz*
*/wd/data/m64077_191108_195203.Q20.fasta.gz*
